# Inducible genetic ablation of *Immt* induces a lethal disruption of the MICOS complex

**DOI:** 10.1101/2023.08.22.554261

**Authors:** Stephanie M. Rockfield, Meghan E. Turnis, Ricardo Rodriguez-Enriquez, Madhavi Bathina, Seng Kah Ng, Stephane Pelletier, Peter Vogel, Joseph T. Opferman

## Abstract

The mitochondrial contact site and cristae organizing system (MICOS) is important for cristae junctions (CJ) formation and for maintaining inner mitochondrial membrane (IMM) architecture. As the largest member, MIC60 is the primary scaffold protein for this complex. While MIC60 has been well studied in yeast and cell culture models, its function in mammals is poorly understood. To address this, we developed a mouse model conditionally deleting *Immt* (which encodes MIC60) and found that global *Immt* deletion disrupted the MICOS complex and resulted in lethality within 9 days of tamoxifen treatment. Pathologically, these mice display intestinal defects consistent with paralytic ileus, resulting in dehydration. We also identified bone marrow hypocellularity in tamoxifen-treated mice. However, bone marrow transplants from *Immt*^WT^ mice failed to rescue survival. Altogether, this novel mouse model demonstrates the importance of MIC60 *in vivo*, in both hematopoietic and non-hematopoietic tissues, and provides a valuable resource for future mechanistic investigations into the MICOS complex. Such investigations could include an *in vivo* structure-function analysis of MIC60 functional domains, with characterizations that are relevant to human diseases.

## Introduction

Mitochondria are responsible for many aspects of cellular biology, including ATP synthesis, lipid synthesis, and mediating apoptosis (1). Mitochondrial structures include the outer mitochondrial membrane (OMM), the inner mitochondrial membrane (IMM), which folds into cristae (2, 3), and the matrix, the region within the IMM where mitochondrial DNA is stored (2). Cristae junctions (CJ) house the large multi-protein complex known as the mitochondrial contact site and cristae organizing system (MICOS) (4). The MICOS complex, of which MIC60 (gene name *Immt*) serves as the main scaffold, is important for the formation and morphology of the mitochondrial cristae. MICOS forms a bridge (known as the mitochondrial intermembrane bridge, or MIB) across the intermembrane space (IMS) to the sorting and assembly machinery (SAM) complex (4, 5). The SAM complex resides within the OMM, where it coordinates with the transfer of outer membrane (TOM) complex to translocate proteins into the mitochondria as well as integrating large β-barrel proteins within the outer membrane (6).

Multiple studies have characterized disrupted cristae morphology and reduced mitochondrial respiration following MIC60 expression loss (7, 8, 9, 10, 11, 12); however, these studies were all completed using yeast models or mammalian cell culture models via siRNA-mediated knockdown. To date, the *in vivo* impact of MIC60 loss has not been documented. Therefore, we generated an inducible *Immt* knockout mouse model to determine the biological effects of losing functional MIC60 protein in adult mice. Following tamoxifen treatment, loss of MIC60 expression was confirmed to correlate with reduced MICOS/MIB protein expression, indicating a collapse of the MICOS complex. *Immt* deletion *in vivo* resulted in a swift reduction in mouse survival. These results demonstrate the importance of MIC60 function *in vivo* and highlights a novel model system for future research.

## Results and Discussion

### Inducible, widespread genetic *Immt* deletion results in the rapid death of mice

We first assessed the expression of MIC60 protein across multiple murine tissue types from adult mice (**Fig 1A**). MIC60 protein expression was highest in mitochondria-rich tissues such as heart, liver, kidney, and skeletal muscle (13). We also observed the presence of multiple isoforms, especially in the kidney, which are reportedly generated through alternative splicing (14). Next, we generated a conditional knockout mouse line for *Immt* by introducing intronic loxP sites on either side of exon 3 (present in all murine *Immt* isoforms), then backcrossed to C57BL/6J mice containing ROSA-CreER^T2^ for tamoxifen-inducible expression of Cre recombinase (**Fig 1B**) (15). The deletion of exon 3 is expected to retain the first 40 amino acids from exons 1 and 2, but then cause a frameshift that codes for 5 alternate amino acids before resulting in a premature stop codon before the predicted transmembrane domain. To determine how quickly to anticipate that MIC60 protein could be eliminated after tamoxifen administration, we sought to determine the half-life of MIC60 and other mitochondrial proteins. Using a cycloheximide time course in murine embryonic fibroblasts (MEFs) in the presence of a pan-caspase inhibitor (Q-VD-OPh) to inhibit apoptosis, we found no appreciable decline in MIC60 or MIC10 protein levels up to 24 hours post-treatment (**Fig 1C**). In contrast, the expression of anti-apoptotic MCL-1 was rapidly lost following translation inhibition in MEFs (16). This indicates that MIC60 is a long-lived protein in MEFs. Furthermore, this long half-life of MIC60 is consistent with other studies which demonstrated the long half-life of MIC60 protein in cardiomyocytes, human neurons, and differentiated murine myotube cells (17, 18).

**Figure 1.**
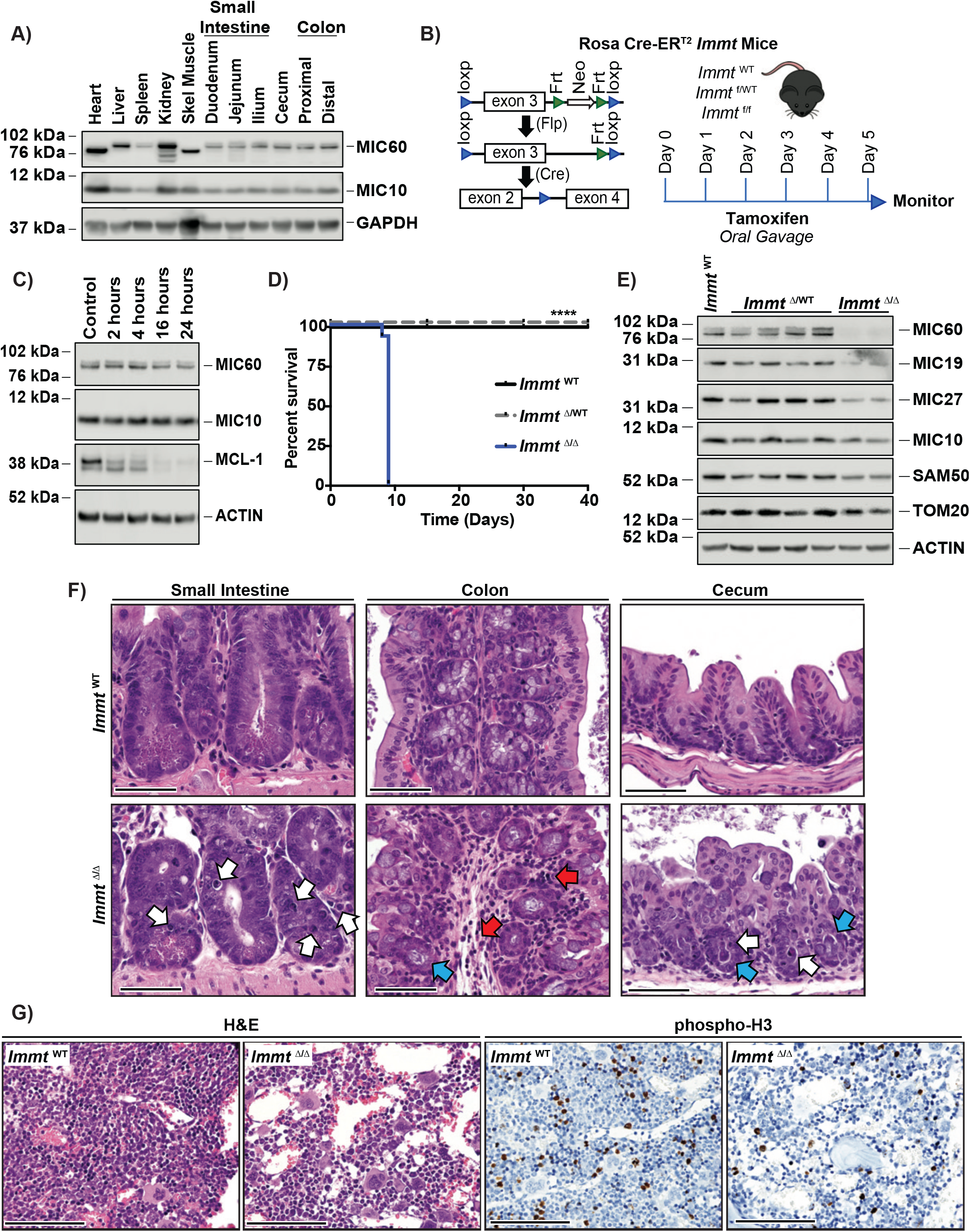
Inducible widespread genetic *Immt* deletion results in the rapid death of mice. **A)** Western blot demonstrating the expression of MIC60 and MIC10, relative to GAPDH, across a variety of murine tissues. Data represents 3 independent experiments. **B)** Diagram depicting the placement of loxp sites to generate tamoxifen-inducible *Immt* deletion in mice and experimental procedure. **C)** Western blot demonstrating the change in mitochondrial protein expression over time following treatment with 10μg/ml CHX (with 40µM Q-VD-OPh) in MEFs. Data represents 3 independent experiments. **D)** Kaplan-Meier survival curve of tamoxifen treatment in *Immt*^WT^ (n=7), *Immt*^f/WT^ (n=12), and *Immt*^f/f^ (n=14). The Log-rank Mantel-Cox test was used to assess significance, p<0.0001. **E)** Western blot from small intestines 9 days after tamoxifen treatment, demonstrating the expression of MICOS and mitochondrial proteins across genotypes. Data represents at least 3 independent experiments. **F)** Representative images of H&E-stained small intestine (left), colon (center), or cecum (right) tissues 7 days after tamoxifen treatment. White arrows indicate apoptotic cells, red arrows indicate submucosal inflammation and edema, and blue arrows indicate dilated crypts with bacteria. Images are at 40X magnification. **G)** Representative images of H&E (left) or phosphorylated Histone 3 (right) stained bone marrow 7 days after tamoxifen treatment. Images are at 40X magnification.

Tamoxifen was provided via oral gavage over a course of 5 days. Whereas *Immt*^WT^ or *Immt*^Δ/WT^ mice were unaffected, *Immt*^Δ/Δ^ mice failed to survive more than 9 days past the initiation of tamoxifen treatment (**Fig 1D**). When *Immt*^Δ/Δ^ mice approached a humane endpoint (based on visual assessment of hunched appearance, minimal movement, and bloated abdomen) (19), they presented with enlarged stomachs and fluid retention in their intestines, cecum, and colon (**Fig EV1A**). Based on this observation, we first focused our analysis on the intestinal tracts of these animals. Western blot analysis of the small intestines (**Fig 1E**) and colon (**Fig EV1B**) of tamoxifen-treated mice revealed that MIC60 protein levels were reduced, and this corresponded with a reduced protein expression of other MICOS and MIB complex members (MIC19, MIC27, MIC10, and SAM50), as described previously (10, 11, 12). Histological analysis of the intestinal tract (small intestines, colon, and cecum) revealed apoptotic cells (white arrows), vacuolar degeneration of small intestine enterocytes, dilated intestinal crypts containing bacteria (blue arrows), and submucosal inflammation (red arrows, **Fig 1F**). These gross findings and histopathology are consistent with a diagnosis of paralytic ileus (20), malabsorption, and bacterial overgrowth. The combined effects of a compromised intestinal mucosa, small intestinal bacterial overgrowth, and paralytic ileus result in decreased water absorption and endotoxemia, which ultimately lead to severe dehydration and death. We also identified reduced expression of MIC60 and MIC10 in the spleen of *Immt*-deleted mice (**Fig EV1C**). Notably, MIC60 expression was reduced but not completely ablated in liver tissue (**Fig EV1D**), and not reduced at all in kidney (**Fig EV1E**), heart (**Fig EV1F**) or skeletal muscle (**Fig EV1G**), suggesting that these mice may be succumbing to the effects of *Immt* deletion in the digestive tract before efficient deletion occurred in other tissues.

Aside from the above-mentioned intestinal aberrations, the only other pathological finding following genetic *Immt* deletion was cellular hypoplasia within the bone marrow (BM) that was not observed in *Immt*^WT^ mice, as revealed by hematoxylin and eosin (H&E) stains of murine femurs (**Fig 1G**). Reduced bone marrow cellularity corresponded with reduced expression of phosphorylated histone-3 (phospho-H3) as a marker of cellular proliferation (0.64% positive cells per µm^2^ in *Immt*^Δ/Δ^ versus 3.9% positive cells per µm^2^ in *Immt*^WT^, **Fig 1G**), suggesting that inducible *Immt* deletion may trigger BM failure. To test this hypothesis, *Immt*^WT^ and *Immt*^f/f^ mice were lethally irradiated to deplete the BM cells and then reconstituted with donor marrow from either *Immt*^WT^ or *Immt*^f/f^ ROSA-CreER^T2^ mice, with subsequent treatment with tamoxifen after engraftment (**Fig 2A**). As shown in **Fig 2B**, transplanting *Immt*^f/f^ BM into *Immt*^WT^ mice improved survival of tamoxifen-induced deletion to two weeks (blue line), which corresponds with the typical time course of BM failure after lethal irradiation (21). However, transplanting *Immt*^WT^ BM into *Immt*^f/f^ failed to rescue, and markedly worsened survival relative to *Immt* deletion (dotted red line, compare to survival curve in **Fig 1D)**. Representative H&E images of the BM following transplants depicts exacerbated hypocellularity after tamoxifen treatment in *Immt*^f/f^ transplanted into *Immt*^WT^ mice relative to *Immt*^WT^ controls, although the BM from *Immt*^WT^ transplanted into *Immt*^f/f^ remained normal (**Fig 2C**).

**Figure 2.**
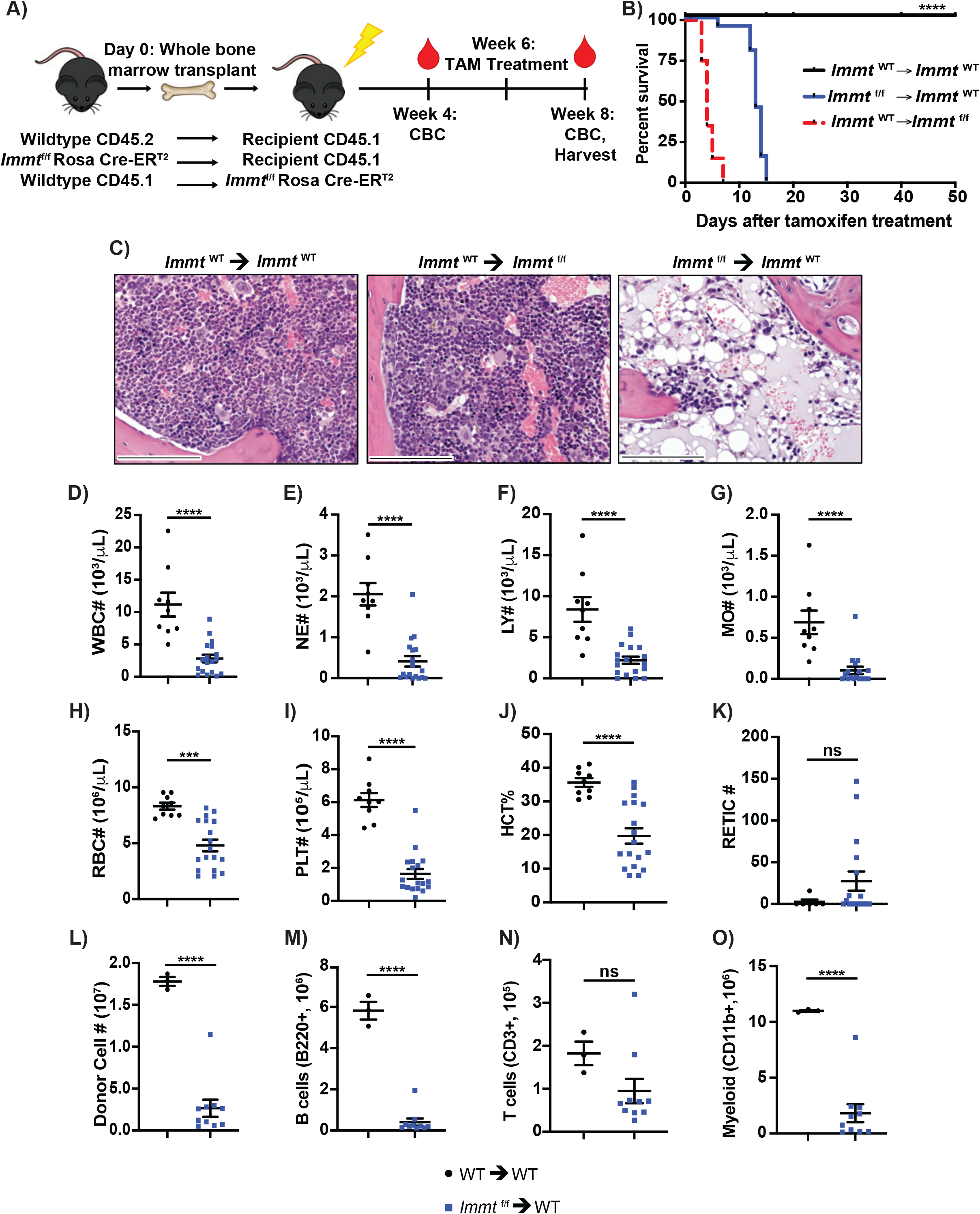
Hematopoietic *Immt*-deletion leads to bone marrow failure. **A)** Diagram summarizing the experimental procedure. **B)** Kaplan-Meier survival curve of tamoxifen treatment in *Immt*^WT^ (CD45.2^+^) transplanted into *Immt*^WT^ (CD45.1^+^, n=16), *Immt*^WT^ (CD45.1^+^) transplanted into *Immt*^f/f^ (ROSA-CreER^T2^ (CD45.2^+^), n=20), and *Immt*^f/f^ (ROSA-CreER^T2^ (CD45.2^+^) transplanted into *Immt*^WT^ (CD45.1^+^, n=21). The Log-rank Mantel-Cox test was used to assess significance, p<0.0001. **C)** Representative images of H&E-stained bone marrow from the indicated transplant experiments, following tamoxifen treatment. All histology images are at 40X magnification. Dot plots (where each dot represents an individual mouse) depicting circulating blood cell (CBC) counts (**D-K)** and donor bone marrow cells (**L-O**); comparisons were made between wildtype controls (black circles) and *Immt*^f/f^ (ROSA-CreER^T2^, CD45.2^+^) transplanted into *Immt*^WT^ (blue squares) after tamoxifen treatment. **D**) White blood cells; **E**) Neutrophils; **F**) Lymphoblasts; **G**) Monocytes; **H**) Red blood cells; **I**) Platelets; **J**) Hematocrit percent; **K**) Reticulocytes; **L**) Cell counts; **M**) B cells; **N**) T cells; and **O**) Myeloid cells. Error bars represent standard error of the mean.

### Hematopoietic *Immt*-deletion leads to bone marrow failure

We next analyzed the circulating blood cell populations by CBC from *Immt*^WT^ BM transplanted into *Immt*^WT^ and *Immt*^f/f^ transplanted into *Immt*^WT^ mice after tamoxifen treatment (**Fig 2D-K**). Additionally, at the conclusion of the experiment, or when mice became moribund, BM was harvested and analyzed by flow cytometry (**Fig 2L-O**). All *Immt*-deleted BM parameters observed, except for reticulocyte (RETIC) numbers and T cell numbers, were significantly decreased compared to the wildtype controls. While T cell numbers trended towards being reduced in *Immt*^f/f^ transplanted into *Immt*^WT^ (relative to controls, **Fig 2N**), the RETIC numbers trended towards being increased in *Immt*^f/f^ transplanted mice (**Fig 2K**), corresponding with the significantly reduced red blood cell (RBC) numbers observed in **Fig 2H** and suggestive of BM stress (22). All together, these results highlight the importance of functional MIC60 in BM reconstitution.

We additionally assessed the same CBC and bone marrow parameters when transplanting BM from *Immt*^WT^ into *Immt*^f/f^ mice, relative to wildtype controls. Quickly repopulating cells such as neutrophils (NE) and monocytes (MO) can be found in recipient blood within one week of transplant (23). Though NE (**Fig 3B**) and platelet (PLT, **Fig 3F**) numbers were unchanged relative to controls, we observed significantly reduced white blood cell (WBC, **Fig 3A**), lymphocyte (LY, **Fig 3C**), and MO (**Fig 3D**) numbers from *Immt*^WT^ transplanted into *Immt*^f/f^ mice. We also identified significantly reduced donor cells, B cells, T cells, and myeloid cells (**Fig 3I-L**). Intriguingly, RBC numbers (**Fig 3E**), hematocrit percent (HCT, **Fig 3G**), and RETIC numbers (**Fig 3H**) were all significantly increased relative to wildtype controls. These results indicate a functional wildtype graft on its way to complete reconstitution. However, similar to our observations in **Fig 1D**, *Immt* deletion in the *Immt*^WT^ BM transplanted mice still resulted in rapid death in less than 10 days following tamoxifen administration (**Fig 2B**). Indeed, histological analysis of the intestinal track from *Immt*^WT^ BM transplanted into *Immt*^f/f^ mice revealed submucosal vacuolar degeneration of enterocytes (gray arrows), inflammation (black arrows), and sloughing of apoptotic mucosal epithelial cells (yellow arrows) that were not present in the wildtype controls (**Fig 3M**). These results therefore support an improperly functioning *Immt*^Δ^ recipient niche and demonstrate that wildtype donor marrow is unable to rescue the intestinal damage observed following tamoxifen treatment in *Immt*^f/f^ mice.

**Figure 3.**
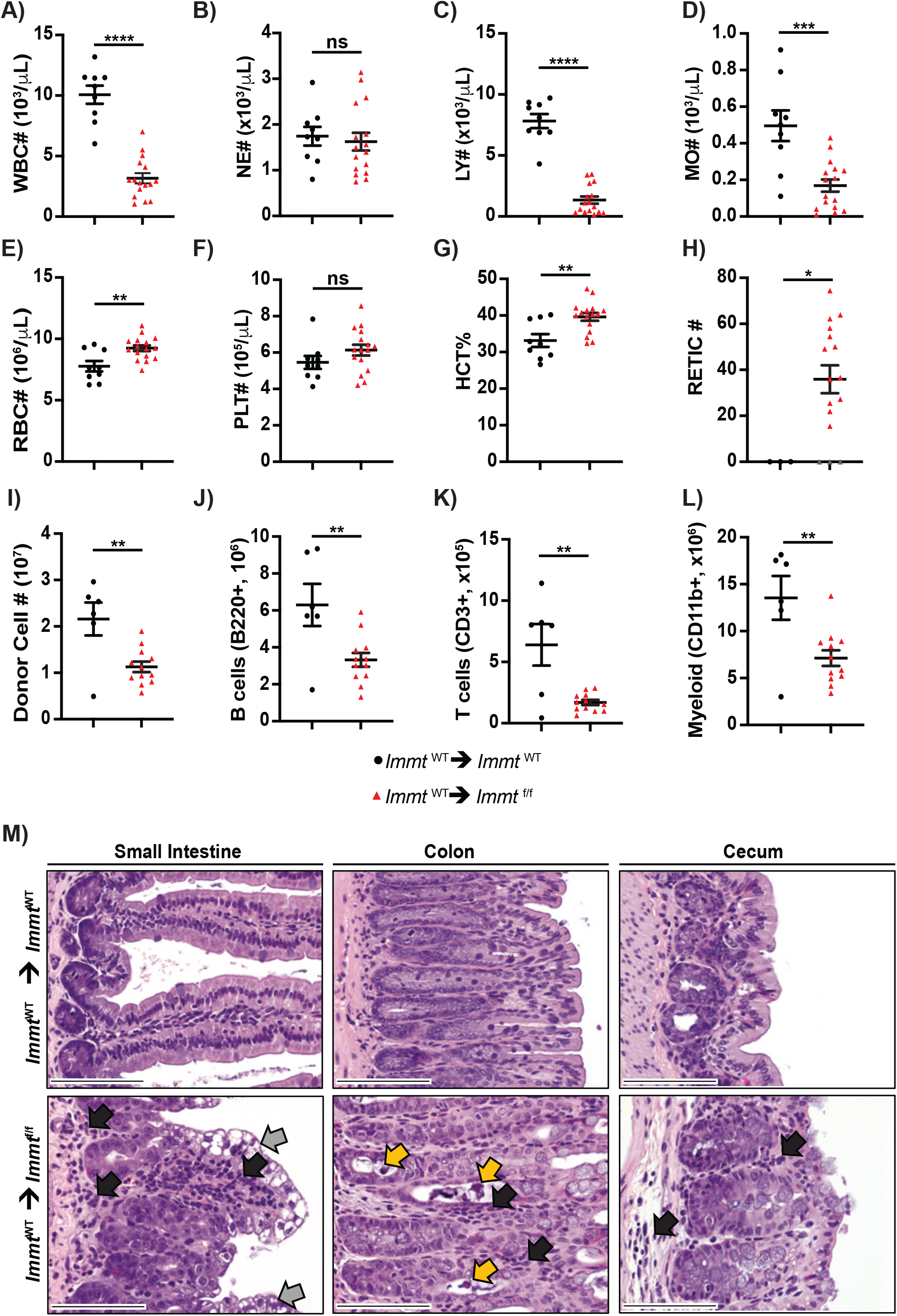
*Immt*^WT^ bone marrow transplantation does not rescue intestinal defects caused by *Immt*-deletion. Dot plots (where each dot represents an individual mouse) depicting CBC counts (**A-H**) and donor bone marrow cells (**I-L**); comparisons were made between wildtype controls (black circles) and *Immt*^WT^ (CD45.1^+^) transplanted into *Immt*^f/f^ (ROSA-CreER^T2^, CD45.2^+^, red triangles) after tamoxifen treatment. **A**) White blood cells; **B**) Neutrophils; **C**) Lymphoblasts; **D**) Monocytes; **E**) Red blood cells; **F**) Platelets; **G**) Hematocrit percent; **H**) Reticulocytes; **I**) Cell counts; **J**) B cells; **K**) T cells; and **L**) Myeloid cells. Error bars represent standard error of the mean. **M**) Representative images of H&E-stained small intestine (left), colon (center), or cecum (right) tissues after tamoxifen treatment of the indicated bone marrow transplanted mice. Black arrows indicate inflammatory cell infiltrates, gray arrows indicate vacuolar degeneration of enterocytes, and yellow arrows indicate sloughing of apoptotic mucosal epithelial cells. Images are at 40X magnification.

Herein, we demonstrate the biological importance of MIC60 through whole-body deletion of *Immt*. To our knowledge, this is the first report to investigate the effects of ablating *Immt*/MIC60 *in vivo*. Importantly, we demonstrate that MIC60 loss corresponds with reduced expression of other MICOS and MIB protein members and thus confirm its role *in vivo* as a scaffold for these complexes. This *in vivo* system provides a means of investigating the importance of different MIC60 structural domains and how MIC60 interacts with other proteins and is regulated in mammals. To date, a small number of disorders have been directly linked to MIC60 dysfunction. PINK1 (Parkinson’s disease (PD)-linked Ser/Thr kinase) was found to phosphorylate MIC60, and rare mutations were identified in Parkinson’s disease patients which associated with impaired locomotive movement in flies (24). More recently, a potentially causal mutation in *IMMT* was identified in two consanguineous patients presenting with developmental encephalopathy (25), and low expression of MIC60 correlated with poor patient prognosis for pancreatic ductal adenocarcinoma (26). Thus, our conditional *Immt* knock-out mouse model provides a valuable resource for future investigations of MIC60 dysfunction *in vivo*.

## Materials and Methods

### Generation of *Immt* conditional knockout mouse

BAC clones (RP24-370K1, RP24-295O10, and RP24-268L22) were purchased from the Children’s Hospital Oakland Research Institute (CHORI) BACPAC Resources (Emeryville, CA). **EV Table 1** summarizes all sequencing primers utilized to identify the BAC clone containing murine *Immt*, transfer the sequence to pBR322 (grown and selected in EL350 cells), introduce a Neomycin cassette downstream of exon 3 (surrounded by Frt flippase recognition targets), and introduce loxP sites on either side of exon 3 (**Fig 1B**; exon 3 is present in all known isoforms of *Immt*, see NCBI GeneID 76614, NC000072.7, Reference GRCm39 C57BL/6J). The constructed plasmid was electroporated into murine embryonic stem cells, then injected into early mouse embryos and implanted as previously described (27). Resulting chimeras were then backcrossed to C57BL/6J mice with ROSA-CreER^T2^ (strain #008463, Jackson Laboratories, Bar Harbor, ME, USA). The final genotypes of the utilized mice were *Immt*^WT^, *Immt*^f/WT^, or *Immt*^f/f^, all with two copies of ROSA-CreER^T2^. Mice were kept on a 12 hour light/dark cycle with food and water provided *ad libitum*; all procedures were approved by the St Jude Institutional Animal Care and Use Committee.

Tamoxifen (T5648, Sigma Aldrich, St. Louis, MO, USA) was prepared in 100% Sunflower Oil (88921, Sigma Aldrich). 1 mg/mouse was fed via oral gavage every day for 5 days, after which the mice were monitored for health and survival. For blood chemistry analysis, ∼50-70 µl of retro-orbital blood was collected and red blood cells were lysed prior to staining with the following BioLegend antibodies for flow cytometry analysis: CD3 (clone 145-2C11), B220 (clone RA3-6B2), GR1 (clone RB6-8C5), and CD11b (clone M1/70).

### Bone marrow transplants

Transplant studies were completed as previously described (28). Briefly, *Immt*^WT^ (CD45.1^+^) or *Immt*^f/f^ (ROSA-CreER^T2^, CD45.2^+^) donor bone marrow was transplanted into recipient 1100 rad irradiated *Immt*^WT^ (CD45.1^+^) or *Immt*^f/f^ (ROSA-CreER^T2^, CD45.2^+^) mice such that the 3 experimental groups were *Immt*^WT^ (CD45.2^+^) into *Immt*^WT^ (CD45.1^+^), *Immt*^f/f^ (ROSA-CreER^T2^, CD45.2^+^) into *Immt*^WT^ (CD45.1^+^), and *Immt*^WT^ (CD45.1^+^) into *Immt*^f/f^ (ROSA-CreER^T2^, CD45.2^+^). After 4 weeks, blood samples were collected to verify engraftment prior to tamoxifen treatment (as detailed above) to delete floxed *Immt*. Murine health was assessed and when mice reached a humane endpoint, blood samples were again collected and analyzed via flow cytometry.

### Immunohistochemical and pathological studies

When tamoxifen-treated mice reached a defined humane endpoint, they were euthanized via CO_2_ and tissues were collected for analyses. For immunohistochemistry, tissue samples were excised, rinsed in PBS, and fixed in 4% paraformaldehyde. All samples were provided to the Comparative Pathology Core (St. Jude Children’s Research Hospital, Memphis, TN, USA) for paraffin embedding, slicing, and staining with Hematoxylin & Eosin or anti-phosphorylated Histone 3 (H3) antibody (Bethyl Laboratories, IHC-00061, 1:200). For all experiments, slides were independently analyzed by a board-certified pathologist.

### Cell Culture

Wildtype murine embryonic fibroblasts (MEFs) were seeded into 6-well tissue culture plates at 15,000 cells/cm^2^. Cells were treated with 10µg/ml cycloheximide (Sigma) and 40µM Q-VD-OPh (Sigma) for the indicated timepoints before harvesting proteins for western analysis.

### Immunoblotting

Tissue fragments were flash frozen in liquid nitrogen upon collecting, then broken apart with a tissue pulverizer (Bessman). Tissue powder or cell pellets were lysed in RIPA buffer for 30min on ice. Protein levels from tissues and cells were assessed as previously described (16). Membranes were horizontally cut to assess proteins of varying molecular weights and incubated overnight at 4°C in freshly prepared primary antibodies diluted in 5% BSA-PBS-T. Antibodies utilized are: MIC60 (Bethyl Laboratories), MIC19 (Invitrogen), MIC27 (Novus Biologicals), MIC10 (GeneTex), SAM50 (Abcam), TOM20 (Santa Cruz), MCL-1 (Rockland), Actin (Millipore Sigma), and GAPDH (Cell Signaling Technologies). Chemiluminescence was read on a Li-COR Odyssey Fc Imager (Li-COR Biosciences, Lincoln, NE, USA) using ImageStudio. No changes were made to image brightness or contrast after acquisition, and if required blots were adjusted for horizontal alignment using ImageJ (NIH, Bethesda, MD, USA).

### Statistical Analyses

All graphs were prepared in GraphPad Prism (Version 9.5.1(528), San Diego, CA, USA). For murine survival studies, a Log-rank Mantel-Cox test, followed by a Gehan, Breslow, Wilcoxon test were completed. For circulating blood cell populations, a standard non-parametric students *t*-test was completed. Statistical significance was set to p≤0.05 (*), p≤0.01 (**), p≤0.001 (***), and p≤0.0001 (****).

## Acknowledgements

This work was supported by the American Lebanese Syrian Associated Charities of St Jude. In addition, the authors would like to thank the members of the Opferman laboratory (St. Jude Children’s Research Hospital) for helpful discussions. The authors also thank the St Jude Children’s Research Hospital Animal Resource Center, Comparative Pathology Core, as well as the Flow Cytometry and Cell Sorting Shared Resource for support of this project.

## Authorship Contributions

J.T.O., S.M.R., M.E.T., R.R.E., and M.B. conceived the study. S.M.R., M.T., R.R.E., and M.B. completed the experiments. J.T.O., S.M.R., and M.E.T. wrote the manuscript. S.M.R. and M.E.T. analyzed the data and prepared figures. S.K.N. and S.P. generated the *Immt* ROSA-CreER^T2^ mouse strain. P.V. performed pathology imaging and analyses. J.T.O. supervised the project.

## Figure Legends

**Figure EV1.**
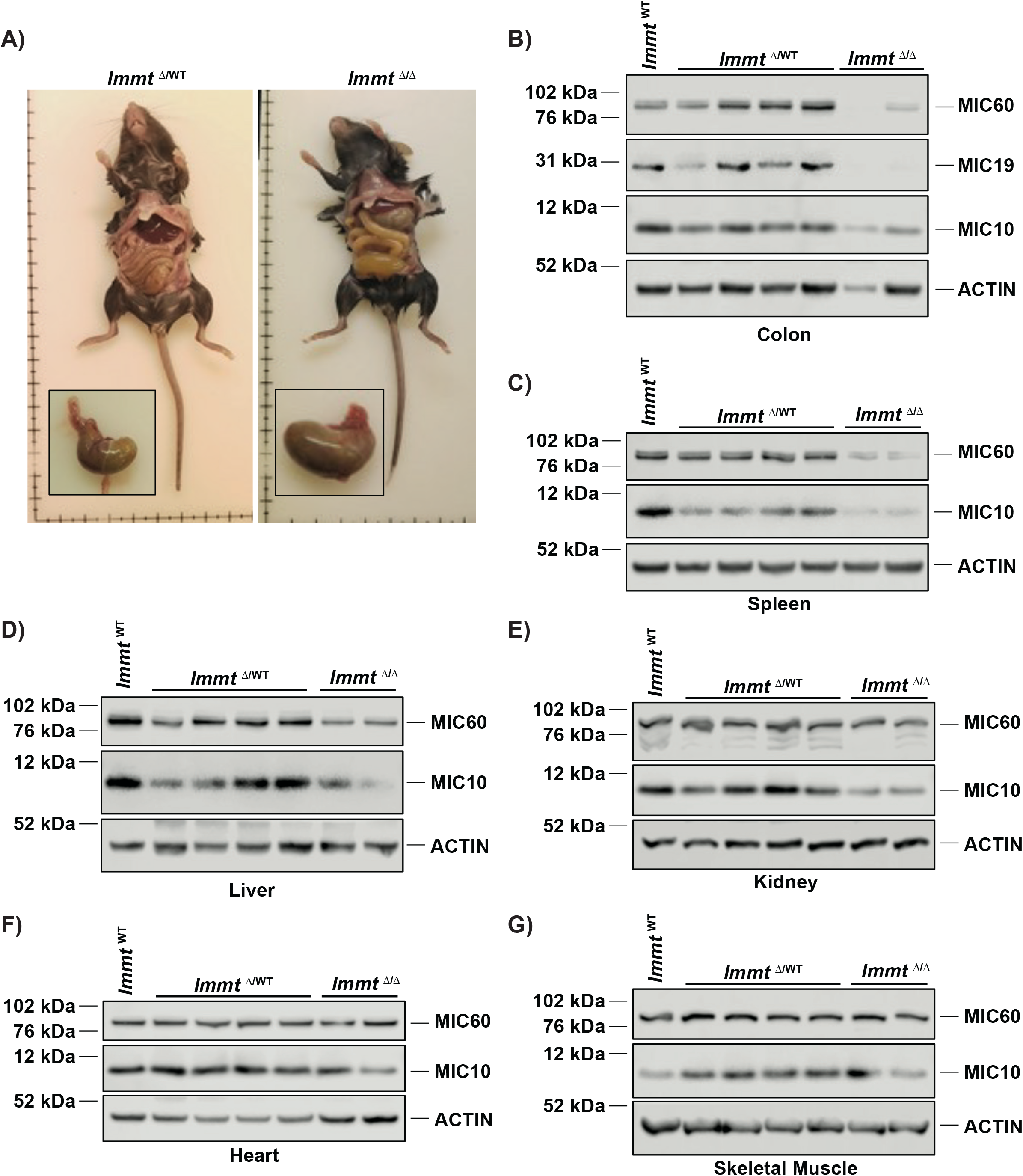
Inducible widespread *Immt* deletion results in intestinal tract defects. **A)** Representative images of murine digestive system following 9 days of tamoxifen treatment. Western blot from **B)** colon, **C)** spleen, **D)** liver, **E)** kidney, **F)** heart, and **G)** skeletal muscle 9 days after tamoxifen treatment, demonstrating the expression of MICOS and mitochondrial proteins across genotypes. Data represents at least 3 independent experiments.

**Extended View Table 1.**
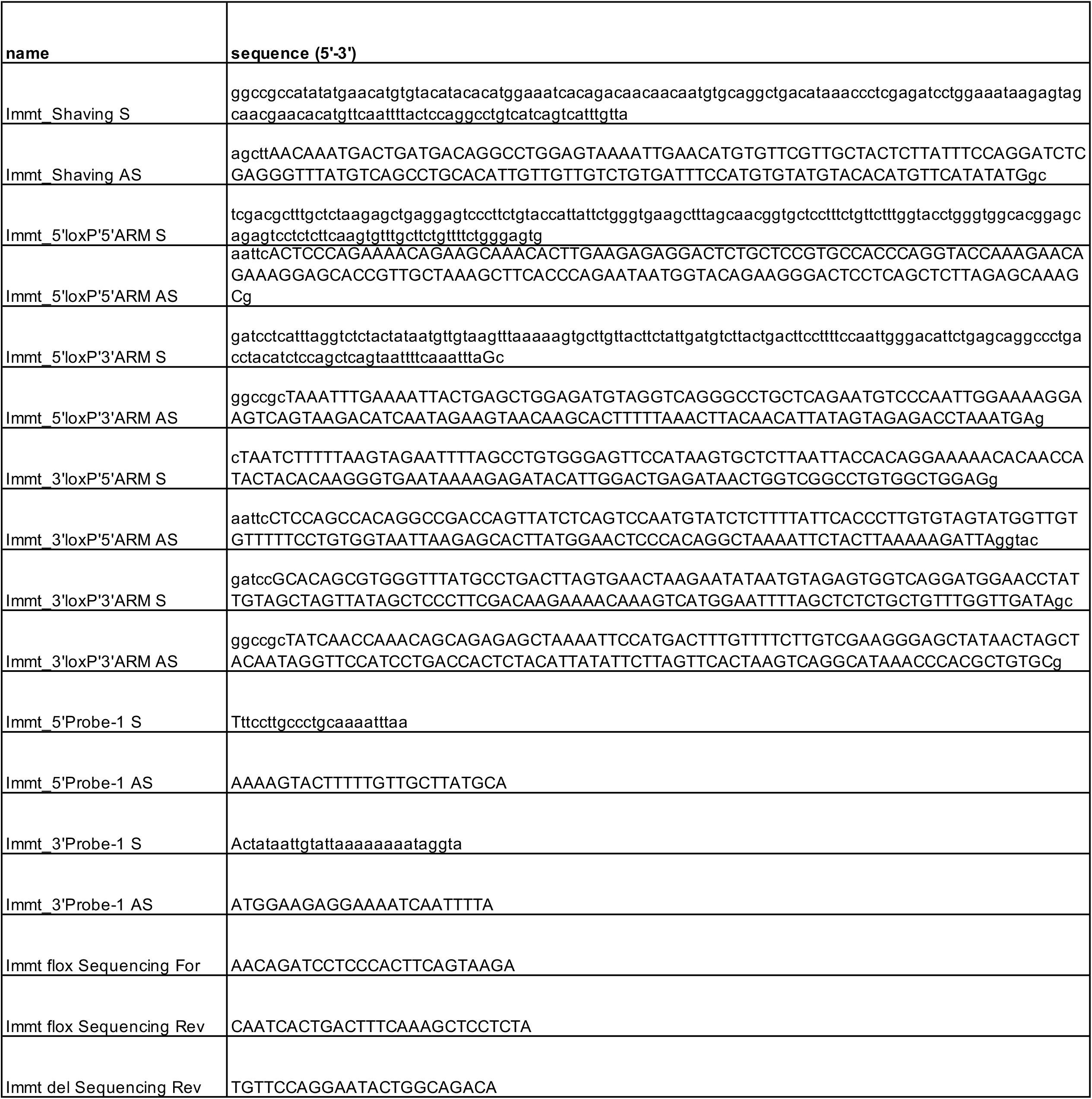

